# Mechanisms that promote the evolution of cross-reactive antibodies upon vaccination with designed influenza immunogens

**DOI:** 10.1101/2022.12.02.518838

**Authors:** Leerang Yang, Timothy M. Caradonna, Aaron G. Schmidt, Arup K. Chakraborty

## Abstract

**SUMMARY:** Immunogens that elicit broadly neutralizing antibodies targeting the conserved receptor-binding site (RBS) on influenza hemagglutinin (HA) may serve as a universal influenza vaccine candidate. Here, we developed a computational model to interrogate antibody evolution by affinity maturation after immunization with two types of immunogens: a chimeric heterotrimeric ‘HAtCh’ antigen that is enriched for the RBS epitope relative to other B cell epitopes, and a cocktail composed of three non-epitope-enriched homotrimeric antigens that comprise the HAtCh. Experiments in mice (Caradonna et al.) find that the chimeric antigen outperforms the cocktail for eliciting RBS-directed antibodies. We show that this result follows from an interplay between how B cells engage these antigens and interact with diverse T helper cells, and requires T cell-mediated selection of germinal center B cells to be a stringent constraint. Our results shed new light on antibody evolution, and highlight how immunogen design and T cells modulate vaccination outcomes.

## INTRODUCTION

The mutability of viruses like human immune deficiency virus (HIV) and influenza poses a major public health challenge. No effective vaccine is available for HIV, and seasonal variation of influenza requires annual vaccine reformulation. Additionally, the severe acute respiratory syndrome coronavirus 2 (SARS-CoV-2) is rapidly evolving variants that reduce the efficacy of current vaccines, thus raising the possibility that booster shots may be required periodically (Cao et al., 2021; Zhou et al., 2021). Developing vaccines that can induce broadly neutralizing antibodies (bnAbs) against highly mutable pathogens could address these challenges. BnAbs can neutralize diverse mutant strains by targeting relatively conserved regions on viral surface-exposed proteins. Although bnAbs for both HIV (Burton et al., 1994; Hraber et al., 2014; Simek et al., 2009) and influenza (Corti et al., 2010; Whittle et al., 2011; Wrammert et al., 2011) have been identified, their natural development is typically rare and delayed (Stamatatos et al., 2009; Sui et al., 2011). Therefore, significant efforts are devoted to designing novel immunogens (Jardine et al., 2015; Kanekiyo et al., 2019; Steichen et al., 2019) or vaccination regimens (Escolano et al., 2016; Torrents de la Peña et al., 2018) that may elicit bnAbs with the ultimate goal of creating so-called “universal” vaccines. The complexity of this challenge has also motivated several theoretical and computational studies focused on the mechanisms underlying bnAb evolution (De Boer and Perelson, 2017; Childs et al., 2015; Ganti and Chakraborty, 2021; Luo and Perelson, 2015; Meyer-Hermann, 2019; Murugan et al., 2018; Nourmohammad et al., 2016; Sachdeva et al., 2020; Shaffer et al., 2016; Sprenger et al., 2020; Wang et al., 2015).

Upon natural infection or vaccination, antibodies are elicited through a Darwinian evolutionary process called affinity maturation (Victora and Nussenzweig, 2012). Naive B cells that express a B cell receptor (BCR) with sufficiently high affinity for antigen (e.g., viral protein) can seed germinal centers (GCs). GC B cells multiply and diversify their BCRs through somatic hypermutation, and subsequently interact with the antigen presented on follicular dendritic cells (FDCs). GC B cells internalize varying amounts of antigen based on the binding affinity of their BCRs to the cognate antigen and then display peptides derived from the antigen complexed with class II MHC molecules (pMHC complexes) on their surface (Nowosad et al., 2016). These B cells compete to interact with T helper cells. Productive interactions result in positive selection that leads to proliferation and mutation, while failure to obtain sufficient help signal triggers B cell apoptosis. Many rounds of mutation and selection ensue, resulting in a progressive increase in B cell binding affinity; some B cells differentiate into memory B cells and plasma cells that produce antibodies (Victora et al., 2010).

BnAb evolution is rare upon natural infection for at least two reasons. First, the overall germline precursor frequency of B cells that target conserved epitopes is relatively rare (Jardine et al., 2016). Furthermore, many germline B cells that target highly variable regions on the antigen can co-seed GCs, and can ultimately out-compete rare bnAb precursors during affinity maturation (Pantophlet et al., 2003). Second, the bnAb precursors may acquire “specializing” mutations and lose their breadth of coverage during affinity maturation (Schmidt et al., 2015a; Wang et al., 2015; Wu et al., 2017). Specialization of bnAbs can occur if the BCR binding footprint is larger than the conserved region on the antigen epitope, which is true for both HIV and influenza RBS. In this case, the BCR can develop strong interactions not with the conserved residues but with the variable residues immediately surrounding them. Therefore, an engineered immunogen that can selectively enrich RBS-directed precursors and also guide them to acquire mutations that promote neutralization breadth is necessary for eliciting bnAbs.

Here, we develop a computational model to study the mechanisms that influence the evolution of RBS-directed influenza bnAbs during affinity maturation. Toward this goal, we study the relative efficacy of RBS-directed B cell evolution upon vaccination with two different types of antigens designed by Caradonna et al. (Caradonna et al., 2022). Both immunogens are “resurfaced” hemagglutinin (rsHA) antigens, where the RBS epitope of H1 A/Solomon Islands/03/2006 (H1 SI-06) is grafted onto antigenically distinct H3, H4, and H14 HA head scaffolds (Bajic et al., 2020) (**Fig. 1A)**. The first class of immunogen is an HA trimeric chimera or ‘HAtCh’, a cystine-stabilized rsH3-rsH4-rsH14 heterotrimer each presenting the same H1 SI-06 RBS epitope; due to the antigenic distance between the H3, H4, and H14 scaffolds, the RBS epitope is enriched relative to all other epitopes. The second class is a cocktail of non-epitope-enriched homotrimers of each rsHA; this cocktail contains the same rsHA monomers as the HAtCh but arranged in a set of homotrimers rather than a single heterotrimer.

**Figure 1.**
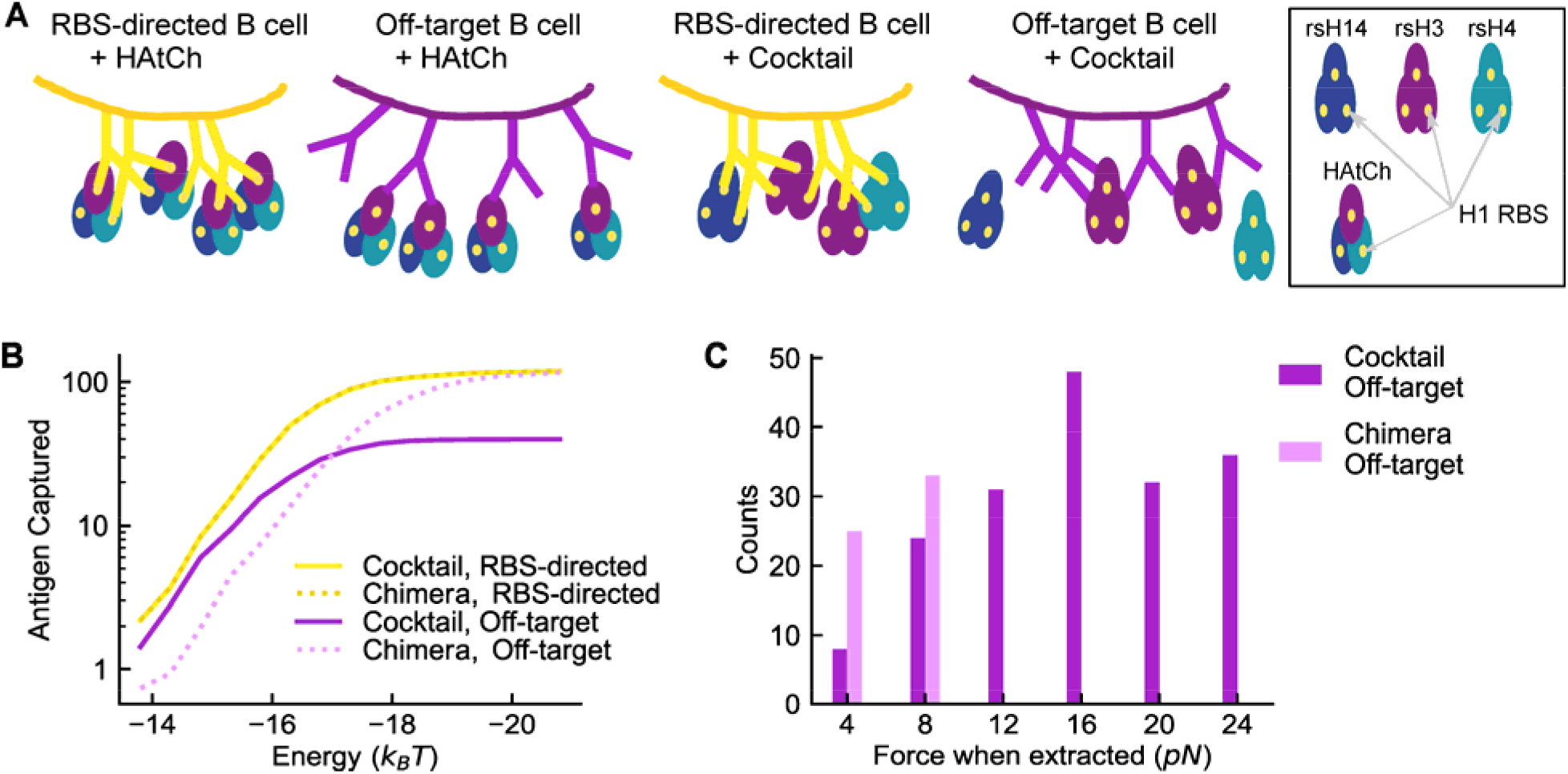
Antigen Capture by B Cells. **(A)** Schematic of the valency of antigens bound by RBS-directed and off-target B cells when interacting with either HAtCh or the rsHA cocktail. **(B)** Amount of antigen captured as a function of binding affinity. For the RBS-directed B cell, the case when the binding affinity towards all three variants is equal is shown. The two antigens are equivalent in this case. **(C)** The forces on FDC-antigen bonds when either the cocktail or the chimeric antigen is captured by strain-specific off-target B cells of low-affinity (-14.8 *k*_*B*_*T*). The histogram was constructed from data from 30 simulations. See also **Figure S1**.

Caradonna et al. report that immunization with the chimeric and cocktail immunogens in mice both elicit cross-reactive RBS-directed B cells, but the chimeric antigen qualitatively outperforms the cocktail. Our computational results reveal the mechanism underlying this result. By studying these new complex antigens, we show how the outcome of GC processes is determined by the interplay of multiple factors: how B cells engage with these immunogens and internalize antigen, the diversity of T helper cells that GC B cells can interact with, and the stringency of T helper cell-mediated selection. We find that upon immunization with a cocktail of homotrimers, only the bnAb precursors can interact with T cells of diverse specificities, while the strain specific B cells must rely on a restricted set of helper T cells. In contrast, upon immunization with the chimeric heterotrimer, both bnAb precursors and strain specific B cells can interact with T cells of diverse specificities. So, intuition may lead us to the conclusion that immunization with the cocktail of homotrimers should perform better than the chimeric heterotrimer at promoting bnAb evolution. The experiments show that the opposite is true. This is because, upon immunization with the chimeric antigen, the bnAb precursors internalize far more antigen than the strain-specific B cells, while these two types of B cells internalize similar amounts of antigen upon immunization with a cocktail of homotrimers. We show that the chimeric antigen performs better as a result of more effective antigen internalization coupled with helper T cells stringently discriminating between B cells based on the amount of pMHC displayed.

Our results also help resolve a controversy in the literature. Gitlin et al. showed that T cell help is a stringent constraint on the selection of GC B cells (Gitlin et al., 2014), while another study suggested that this was not so (Yeh et al., 2018). Our finding that T cell help must be a stringent constraint on B cell evolution in the GC helps resolve this debate. Taken together, our study, and that of Caradonna et al., highlight the importance of immunogen design and T helper cells in determining vaccination outcomes and suggest that modulating these effects is necessary to elicit RBS-directed influenza bnAbs.

## RESULTS

### Model development

#### Seeding the germinal center

We simulate GC reactions induced by either the cocktail of three homotrimeric rsHA antigens or the corresponding epitope-enriched heterotrimeric chimera, ‘HAtCh’ immunogen, described above and in the companion paper (Caradonna et al., 2022). Because off-target germline B cells outnumber the RBS-directed bnAb precursors (Kuraoka et al., 2016; Schmidt et al., 2015a), we seed each GC with 99 strain-specific off-target B cells and 1 RBS-directed bnAb precursor, making the total founder number representative of GCs in mice (Tas et al., 2016). Since the three HA scaffolds are antigenically distinct, an off-target B cell can recognize only one component, which is randomly designated at the beginning of the simulation; mutations change the affinity towards this component. An RBS-directed precursor can target all three components. The initial free energy of binding (or affinity) is set to be *E*_*a*_ for the target component for strain-specific B cells. For simplicity, the RBS-directed precursors are assumed to initially bind all three components with affinity, *E*_*a*_, but as affinity maturation progresses the affinity of an RBS-directed B cell for the three components can become different and even be below the recognition threshold for some components. The amounts of antigen captured by the RBS-directed B cells are determined by all three binding affinities. The absolute value of *E*_*a*_ does not affect the results because all other free energies are scaled to this reference. We choose *E*_*a*_ *= -13*.*8 k*_*B*_*T*, where *k*_*B*_ is the Boltzmann constant and *T* is the temperature (∼300 K), because it corresponds to a dissociation coefficient, *K*_*d*_, of 1 µM, which is approximately the threshold for naive B cell activation (Batista and Neuberger, 1998).

#### Selection, proliferation, and mutation

To model the GC dynamics in mice, founder B cells divide four times without mutation, and then, the competitive phase of affinity maturation lasts for 28 cycles, or ∼14 days (Meyer-Hermann et al., 2012; Victora et al., 2010). Each cycle, B cells that fail positive selection are subsequently removed from the GC via apoptosis. Additionally, ∼10 % of positively selected B cells stochastically differentiate into memory and plasma cells, and exit the GC; the remaining GC B cells proliferate and further mutate.

A positively selected B cell divides twice and one daughter cell can mutate in each division. A mutation is either fatal, silent, or affinity-changing with probabilities of 0.3, 0.5 and 0.2, respectively (Zhang and Shakhnovich, 2010). The PINT database (Kumar and Gromiha, 2006) shows that affinity changes of protein-protein interfaces upon mutations are more likely to decrease than to increase the binding affinity. We describe this data using a shifted log-normal distribution with parameters chosen so that about 5% of the mutations are beneficial (Sprenger et al., 2020; Wang et al., 2015); for an off-target B cell, *i*, the free energy change due to mutation is given by:

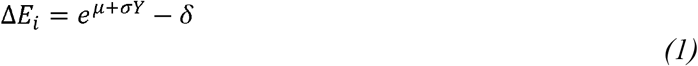

where *Y* is a standard normal random variable, and *µ*, σ, and δ are constants fitted to the empirical distribution (Kumar and Gromiha, 2006).

For RBS-directed B cells, a mutation alters affinities for the individual HA components differently. To model this, we draw three random numbers [*y*_*1*_, *y*_*2*_, *y*_*3*_], one for each component, from a multivariate Gaussian distribution with the mean of zero and the covariance matrix of Λ defined as follows:

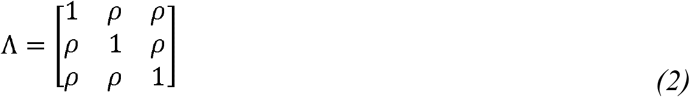

The correlation, ρ, reflects the difference between the antigens and is chosen to be 0.7; we have also carried out calculations with ρ = 0.4. For a given bnAb precursor, *i*, each sampled number, *y*_*j*_, corresponding to the HA component, *j*, is then converted to the free energy change due to mutation, Δ*E*_*ij*_, for this variant using Eq. 1 as follows:

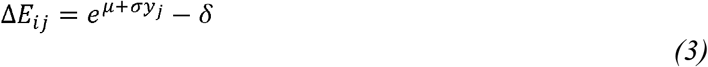

The distribution of free energy changes towards any one component after a mutation is equivalent to that of a strain-specific B cell mutation.

#### Antigen Capture by B Cells

A cross-reactive RBS-directed B cell can bind to a single HAtCh heterotrimer with up to three BCRs, but an off-target B cell can bind a single heterotrimer with, at most, one BCR (**Fig. 1A**). For the cocktail of homotrimers, both an off-target B cell and an RBS-directed B cell can engage a single cognate antigen trimer with multiple BCRs; however, a strain-specific B cell can only recognize a third of the antigen molecules, while a cross-reactive B cell can recognize all the antigen molecules (**Fig 1A**).

GC B cells extract antigens from the surfaces of FDCs using mechanical pulling forces (Nakanski et al., 2013; Nowosad et al., 2016). The B cell synapse interacting with a FDC is modeled as a 2-dimensional circle divided into lattice points occupied by antigen molecules and BCRs (Fleire et al., 2006a; Tsourkas et al., 2007). BCRs and antigen molecules are initially randomly distributed on the lattice. The lattice spacing is 10 nm, which is of the same order as the collision radius of BCR and ligand (Fleire et al., 2006a). During the clustering phase, BCR and antigen molecules diffuse freely and attempt to bind when they are within one lattice point (see STAR METHODS for details). The probability of success is:

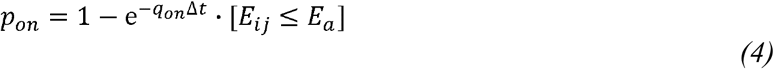

where the Iverson bracket sets the minimum affinity required for binding to be *E*_*a*_, which is equal to the initial precursor affinity, and

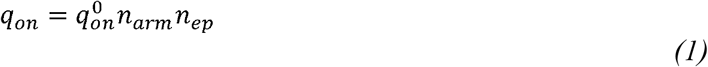

represents the steric factor. This factor is determined by *n*_*arm*_, the number of free BCR arms (between 0 and 2), *n*_*ep*_, the number of free cognate BCR epitopes on the antigen (between 0 and 3), and the basal rate 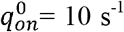. With Δ*t* = 5 × 10^−4^ s, which is the characteristic time scale of diffusion over the lattice, this basal rate results in the successful binding probability of *p*_*on*_ = 5 × 10^−3^. This number approximately accounts for the entropic penalty of aligning two molecules.

An established BCR-antigen bond (labeled, *i* below) breaks with probability,

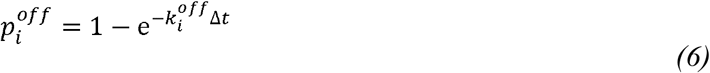

Where 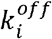is its off-rate. Assuming the activation barrier for bond formation is negligible compared to the binding free energy, the off-rate is related to binding free energy by:

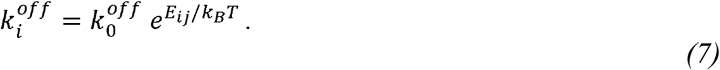

where 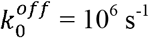 (Batista and Neuberger, 1998), and *E*_*ij*_ is the binding free energy of BCR, *i*, for antigen, *j*.

Our simulations result in the formation of antigen-BCR clusters, followed by antigen internalization through mechanical pulling. We assume that antigen molecules are tethered to the FDC membrane with a binding free energy of -19 *k*_*B*_*T*, which makes antigen capture most sensitive to affinity change in *K*_*d*_ of 1 – 0.01 μM range, but affinity ceiling is reached when *K*_*d*_ < 1 nM (Batista and Neuberger, 1998). A pulling force of 8 pN is applied to each BCR (Nowosad et al., 2016), which is transferred to the antigen-BCR bonds and the FDC-antigen bonds (Amitai et al., 2018). If a BCR is bound to 2 antigen molecules, the force is divided equally by the two arms of the BCR. For a given antigen molecule, the force applied to its FDC-antigen bond is the sum of forces applied by all the BCR arms bound to it. The off-rates of both FDC-antigen and antigen-BCR bonds increase with applied force (Bell, 1978):

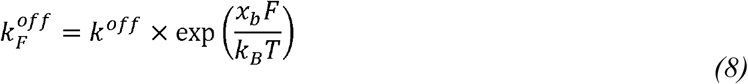

where 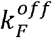 is the off-rate under force, *F* is the force and *x*_*b*_ is the bond length, taken to be 1 nm (Erdmann and Schwarz, 2004). When an antigen-BCR bond breaks, the BCR goes into a refractory state, which prevents instant rebinding with the same antigen (Erdmann and Schwarz, 2004). The duration is taken to be 0.1 s, which is much greater than the antigen diffusion timescale *l*^*2*^*/4D* = 5 × 10^−4^ s. At the end of each time step, any BCR or BCR-antigen cluster that is not connected to the FDC is internalized.

#### Antigen capture depends on antigen design and cross-reactivity of B cells

**Fig. 1B** shows the total amount of antigen captured as a function of BCR binding affinity for cross-reactive and strain-specific B cells for the cocktail and chimeric antigens. Notably, neither immunogen design is better at conferring an advantage to RBS-directed B cells in capturing antigens across the entire affinity range. At low affinity, representative of the early GC, the advantage of cross-reactive B cells over strain-specific B cells is greater for the chimeric antigen while at high affinity, the opposite is true.

At low affinity, antigen availability is not limiting, and the amount of antigen captured is largely determined by the forces imposed on the FDC-antigen bonds by the BCRs bound to the antigen molecules. Multiple off-target BCRs can engage a homotrimeric antigen in the cocktail, but not the chimeric antigen (**Fig. 1A**). Therefore, the forces on the FDC-antigen bonds are typically higher for the homotrimeric antigen bound by strain-specific B cells compared to the chimeric antigen bound by such cells. This point is illustrated quantitatively using results from our simulations. At the low B cell affinity of -14.8 *k*_*B*_*T*, successful extraction of homotrimers in the cocktail frequently results from high forces on FDC-antigen bonds (**Fig. 1C**), enabled by clustering of antigens and BCRs (**Fig. S1**). The maximum possible force of 24 pN is realized when three BCRs are bound to one antigen, each contributing 8 pN of force. Using Eq. 8, the off-rate for the FDC-antigen bond increases by ∼300-fold if an antigen is bound by three BCRs. For the strain-specific B cells capturing chimeric antigen, however, the force on the FDC-antigen bond is always equal to the force on a single antigen-BCR bond because only one BCR can bind to an antigen. Depending upon whether one or both arms of the BCR are bound to an antigen, this force is either 8 pN or 4 pN, respectively. So, the increase in off-rate is relatively modest compared to when strain-specific B cells engage the homotrimeric antigen. This is why strain-specific B cells internalize more of the homotrimeric antigen than the chimeric antigen when antigen-BCR binding affinity is low.

For both types of antigens, cross-reactive RBS-directed B cells can bind an antigen molecule with multiple BCRs. So, at low affinity, these cells capture a larger amount of antigen relative to strain-specific B cells for the chimeric antigen and a similar amount of antigen for the cocktail (**Fig. 1B**).

For high BCR affinity, the cross-reactive B cells capture more antigen than the strain-specific B cells do when interacting with the cocktail of homotrimers (**Fig. 1B**). Beyond a certain affinity, the amount of antigen captured plateaus for the cocktail; this plateau corresponds to the B cell binding affinity approaching the FDC-antigen bond energy of -19 *k*_*B*_*T*. As a result, B cells capture most of the cognate antigens they encounter (**Fig. 1B**). Consequently, antigen availability becomes a limiting factor: cross-reactive B cells are favored because they can bind all antigens while strain-specific B cells only recognize about a third of the antigen molecules presented as a homotrimeric cocktail. For the chimeric antigen, at very high affinity, all antigen molecules can be internalized successfully even with monomeric bonds, so the advantage of cross-reactive B cells is small.

#### RBS-directed B cells evolve more readily upon immunization with chimeric antigen if T cell help is a stringent constraint for positive selection of GC B cells

After antigen capture, B cells compete for positive selection by helper T cells by presenting the T cell epitopes that are derived from the internalized antigen. The homotrimeric cocktail allows only cross-reactive B cells to capture diverse rsHA components, while the chimeric design allows both cross-reactive and strain-specific B cells to internalize all three components (**Fig. 1A**). If the T cell epitopes contained in each rsHA variant are distinct sets, then upon immunization with a cocktail, only the cross-reactive B cells can interact with diverse T cells (**Fig. 2A**). This is because each T cell is specific for its epitope, and a single mutation within a TCR epitope or flanking sites can abrogate recognition (Birnbaum et al., 2014; Carson et al., 1997; Huseby et al., 2005, 2006).

**Figure 2.**
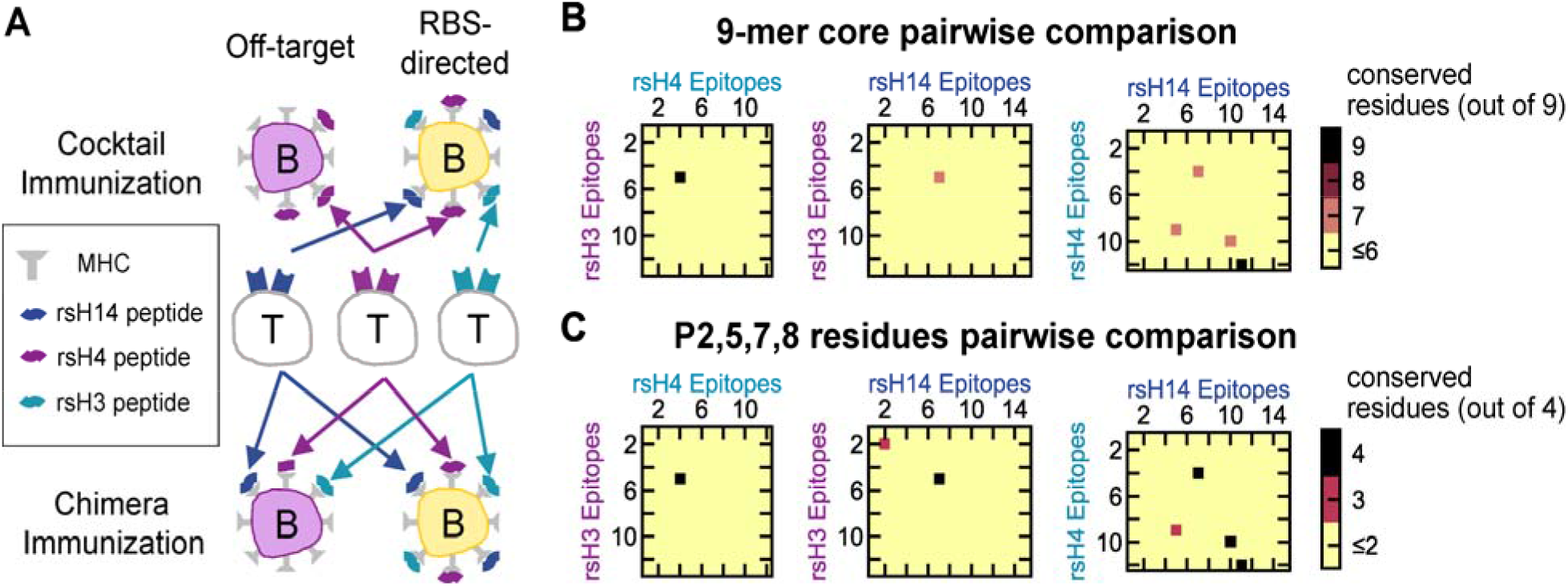
Selection by T Cells. **(A)** Schematics showing the differences between how cross-reactive and strain-specific B cells interact with helper T cells. For the cocktail immunization, only the cross-reactive RBS-directed B cells present pMHCs derived from multiple rsHA components. For the chimera immunization, both strain-specific and cross-reactive B cells present pMHCs from all three rsHA components. **(B-C)** Pairwise comparison of computationally predicted helper T cell epitopes in the rsHA antigens. Each axis corresponds to the ranks of the top 20 percentile predicted 15-mer T cell epitopes, derived from the three rsHA variants. (B) Number of conserved residues in pairwise comparisons focusing on the 9-mer cores of the peptides. (C) Number of conserved residues in pairwise comparisons focusing on the P2, P5, P7 and P8 residues of the 9-mer cores of the peptides. See also **Table S1, Figure S2**.

The rsHA components use antigenically distinct scaffolds derived from different subtypes, which results in the antigenic distances between the overall proteins (except for the RBS epitope) to be very large. The sequence homology between the rsHA components are 58.4% (rsH3-rsH4), 60.5% (rsH3-rsH14), and 72.5% (rsH4-rsH14). The large antigenic distance between the scaffolds raises the possibility that the components in the cocktail carry distinct T cell epitopes.

We used the Immune Epitope Database and Analysis Resource (IEDB) MHCII binding prediction tool to analyze the predicted T cell epitopes in the H3, H4, and H14 rsHA antigens (**Table S1**) (Jensen et al., 2018; Nielsen and Lund, 2009; Nielsen et al., 2007; Wang et al., 2008, 2010). Mice immunized with the cocktail or chimeric antigens were mixed 129/Sv and C57BL/6 mice. Therefore, we used the I-A^b^ MHC allele to ask whether the T cell epitopes contained in the three HA components were distinct. None of the predicted 15-mer peptides that ranked in the top 20 percentile against randomly generated peptides were fully conserved in two different variants. When we relaxed the criteria to just the 9-mer cores, only two pairs were conserved in two different variants (Fig. 2B). We further focused on the identity of just P2, 5, 7, and 8 of the cores, which are the most likely TCR-contacting residues for the I-A^b^ haplotype (Nelson et al., 2015). Still, only five pairs were conserved in all pairwise comparisons (**Fig. 2C**). B cells that capture rabbit serum albumin and human serum albumin (76% sequence homology) do not compete with each other due to mutations in T cell epitopes in mice with I-A^b^ haplotype (Woodruff et al., 2018). For this rabbit and human serum albumin, we found 3 pairs of conserved 9-mer cores and 3 pairs of conserved peptides 2, 5, 7, and 8 in both proteins, which is comparable to the resurfaced HA components (**Fig. S2**). Therefore, we conclude that the components of the cocktail likely contain distinct T cell epitopes. We account for this feature in our simulations by keeping track of which antigen a B cell internalizes and partitioning T helper cells into three distinct groups based on their specificity for epitopes derived from each of the HA variants. The number of T cells in each group is the same.

T cells make numerous short contacts with diverse B cells (Allen et al., 2007). For each contact, there is a small chance of it being a productive encounter, which increases with the amount of peptide presented (Shulman et al., 2014). It is conjectured that positive selection likely requires several productive encounters (Dustin, 2014). Therefore, the amount of help a B cell receives will increase with the number of encounters with cognate T cells, which is determined by the types of pMHC it presents, the number of cognate T cells, and the number of competing B cells. Therefore, we represent the probability of positive selection of a B cell *i* as follows:

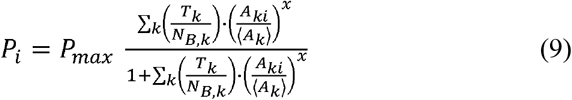

where *A*_*ki*_ is the amount of the HA component *k* internalized by the B cell *i*; *A*_*k*_> is the mean amount of HA component *k* internalized by the B cells that recognize this component; *N*_*B,k*_ is the number of such B cells; and *T*_*k*_ is the number of T cells that target the epitopes from the HA component *k*, which we assume to be equal for all variants. The maximum probability of selection, *P*_*max*_, accounts for the fact that GC B cells are inherently apoptotic irrespective of BCR affinity (Mayer et al., 2017). We can consider *P*_*max*_ to be the chance of avoiding the default fate of apoptosis: 1 – *P*_*apoptosis*_. We chose *P*_*max*_ = 0.6 as it results in good correspondence between the timescales predicted by our model and experiments (Caradonna et al., 2022); other values were also tested, and the qualitative result does not change.

An important feature of the model is the exponent x; larger values of it imply T cell help more stringently depends on the amount of pMHC presented. If x is less than 1, small differences (e.g., 2-fold) in pMHC displayed on two B cells would have a relatively small effect on selection outcome, whereas if x is greater than 1, such small differences would likely lead to the selection of the B cell that displays more antigen.

**Fig. 3A** shows predictions of our model upon immunization with the chimeric and cocktail immunogens for the temporal evolution of the fraction of GC B cells that evolve from RBS-directed bnAb precursors; i.e., B cells that have acquired higher binding affinities than the precursors. A striking feature of these results is that, for immunization with the chimeric immunogen, the evolution of RBS-directed B cells becomes increasingly more efficient as T cell selection becomes more stringent (larger values of *x*); but for immunization with a cocktail of antigens, the opposite is true. **Fig. 3B** graphs a related quantity, the fraction of RBS-directed B cell mutants that can recognize at least two of the three HA components in the immunogens (“scaffold-independent”). A low value indicates that RBS-directed B cells tend to specialize to only one component. Our model predicts that cross-reactive mutants evolve more readily upon immunization with the cocktail when T cell help is permissive, but with the chimera when T cell help is stringent (**Fig. 3B**). The cocktail improves in selecting cross-reactive B cells in late GC if T cell help is stringent (**Fig. 3B**), but by day 14 only a small fraction (12 % for *x* = 1.5) of the simulated GCs still have RBS-directed B cells (**Fig. S3A**). So, our model predicts that cross-reactive RBS-directed B cells will evolve more readily upon immunization with the chimeric antigen, compared to the cocktail, if T cell help is a stringent constraint for positive selection of B cells. **Fig. S3D** and **S3E** show that this qualitative trend is not changed when ρ is changed to 0.4 or when *p*_*max*_ is changed to 1.

**Figure 3.**
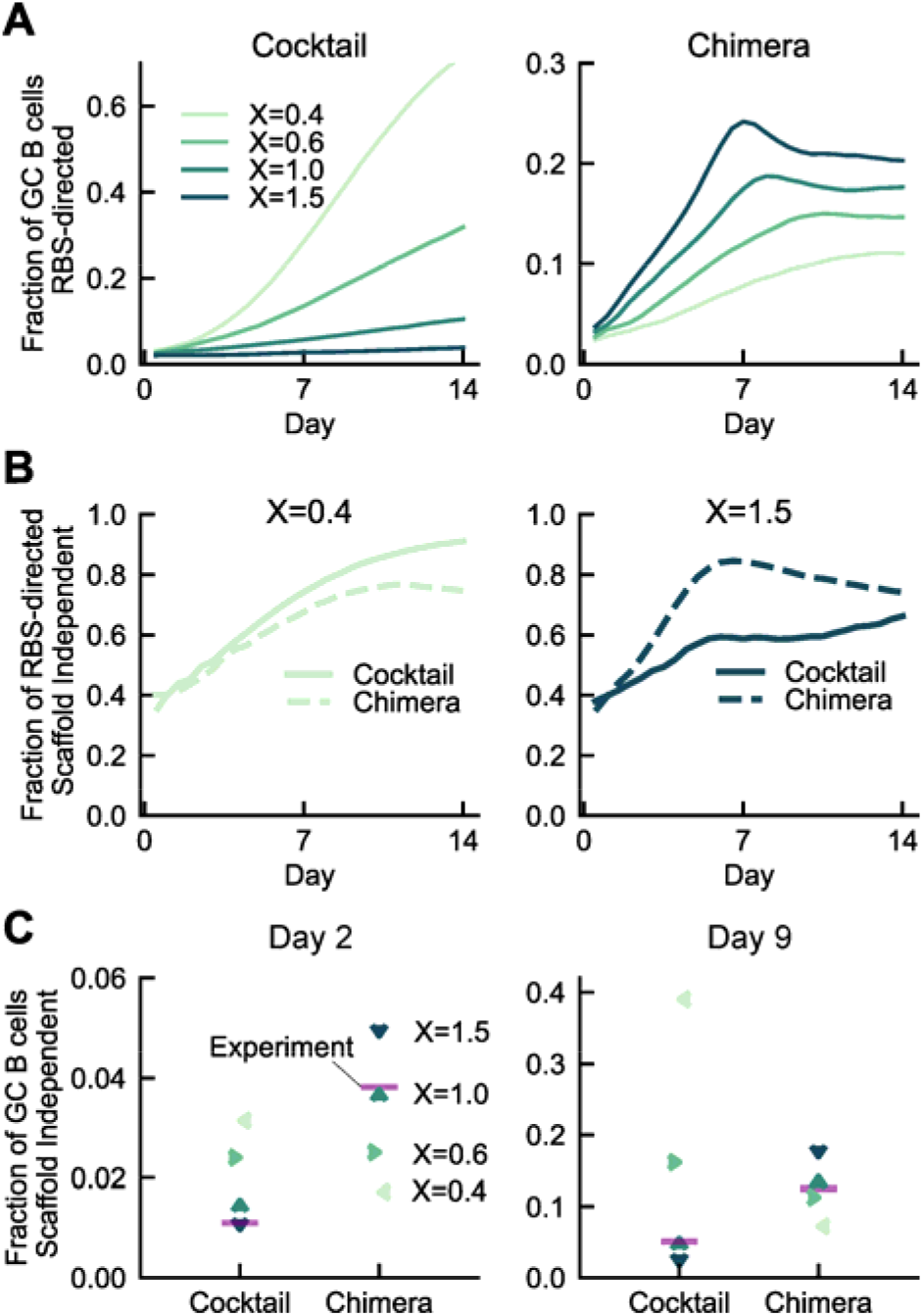
Model predictions and experimental results for the evolution of cross-reactive B cells upon immunization with the cocktail or the chimeric antigen. **(A)** Fraction of GC B cells that are RBS-directed as a function of time in our simulations. Changing the stringency of T cell selection have opposite effects for the chimeric and the cocktail antigens. **(B)** Fraction of RBS-directed B cells that are cross-reactive to at least two rsHA components. When selection by T cells is permissive (*x* < 1), the cocktail favors the evolution of cross-reactive B cells, and the opposite is true for stringent selection (*x* ≥ 1). **(C)** Comparison of model predictions for varying levels of T cell selection stringency with the results of mice immunization experiments at two time points. See also **Figure S3**.

**Fig. 3C** shows the number of RBS-directed GC B cells that bind to at least two of three components as a fraction of all HA-binding GC B cells on days 8 and 15 after mice were immunized with the two types of immunogens (Caradonna et al., 2022). We assume days 8 and 15 post-immunization correspond to days 2 and 9 of GC, since GC initiation typically takes about 6 days (Jacob et al., 1991). While Caradonna et al. report the value as a fraction of all IgG^+^ GC B cells, because our model does not consider background GC B cells that do not bind to any HA, we extract the fraction of RBS-directed B cells among B cells that bind to HA from the experimental data. The qualitative trends in the data are not affected by the background B cells. **Fig. 3C** shows a comparison of our model predictions and experimental data for the fraction of cross-reactive RBS-directed B cells in the GCs on these days. The model predictions are obtained by combining the results in **Figs. 3A** and **3B**.

The experiments show a qualitatively higher frequency of cross-reactive RBS-directed B cells in GCs both day 8 and day 15 after immunization with the chimeric antigen (also see Cardonna et al, 2021). These experimental results are consistent with our predictions when T cell help is stringent but not when it is permissive. The model predicts that, if the T cell help is stringent (*x ≥ 1*), a higher fraction of GC B cells will be RBS-directed and cross-reactive after immunization with the chimeric antigen than with the cocktail (**Fig. 3C**). For example, if *x* = 1, 3.8 % of GC B cells are RBS-directed and cross-reactive on day 2 for the chimeric antigen and 1.5 % for the cocktail. On day 9, the numbers are 14 % for the chimera and 5.5 % for the cocktail. In contrast, if T cell help is permissive (*x* < 1), the cocktail favors the evolution of cross-reactive B cells. For example, if *x* = 0.4, 3.1 % of GC B cells are RBS-directed and cross-reactive on day 2 for the cocktail and 1.7 % for the chimera; the same trend is true at day 9 (39 % for the cocktail and 7.2 % for the chimera). We emphasize that what is important is not the precise numbers noted above, but that the qualitative trend of which type of antigen promotes the evolution of RBS-directed cross-reactive antibodies is opposite for stringent versus permissive selection by T helper cells. The model predictions have the same trend as the experimental data when T cell help is a stringent constraint. Therefore, we conclude that T cell help stringently depends on pMHC density. We also note that even under the most stringent selection tested (*x* = 1.5), stochasticity in evolution allows for clonal heterogeneities inside individual GCs (**Fig. S3B**) (Tas et al., 2016) and broad affinity distribution of B cells both within and across GCs (**Fig. S3C**) (Kuraoka et al., 2016), consistent with previous findings in the literature.

#### Mechanism for why T cell selection stringency promotes cross-reactive B cell evolution for the chimeric immunogen, but not the cocktail

For the chimeric immunogen, cross-reactive RBS-directed B cells can bind to the antigen multivalently, while the off-target B cells cannot, and so the former can capture significantly more antigen than the latter in the early stages of the GC reaction (**Fig. 1**). Events that occur in the early GC are critically important for the RBS-directed precursors as they are few in number and could be easily extinguished due to stochastic effects. Thus, to promote the evolution of RBS-directed B cells, their principal advantage over off-target B cells (more antigen captured) must be amplified by the selection force. This advantage is amplified if positive selection by T helper cells discriminates stringently based on the amount of captured antigen, as this favors the selection of the cross-reactive B cells. Indeed, our simulation results show that the probability that RBS-directed precursors are positively selected in the early GC grows with the value of *x* (**Fig. 4A**) upon immunization with the chimeric antigen. If bnAb precursors are more readily positively selected in the early GC, they multiply more and thus have a higher chance of acquiring the rare mutations that confer breadth. Such an effect of an early advantage affecting future fate has been observed in evolving asexual populations (Nguyen Ba et al., 2019). Consistent with this expectation, our simulation results (**Fig. 4B**) show that upon immunization with the chimeric antigen, mutations that confer breadth are more easily found in the population if T cell selection stringency is high. The resulting cross-reactive cells are then selected for and accumulate because they have a large fitness advantage (**Fig. 4C**, top row). Moreover, although specializing mutations also occur for RBS-directed B cells, they are not selected for because the incurred loss of cross-reactivity would significantly inhibit antigen capture. The advantage of cross-reactive mutations over specializing mutations is pronounced for more stringent selection (**Fig. 4C**, top row). These reasons promote cross-reactive B cell evolution upon immunization with the chimeric antigen if T cell selection is a stringent constraint.

**Figure 4.**
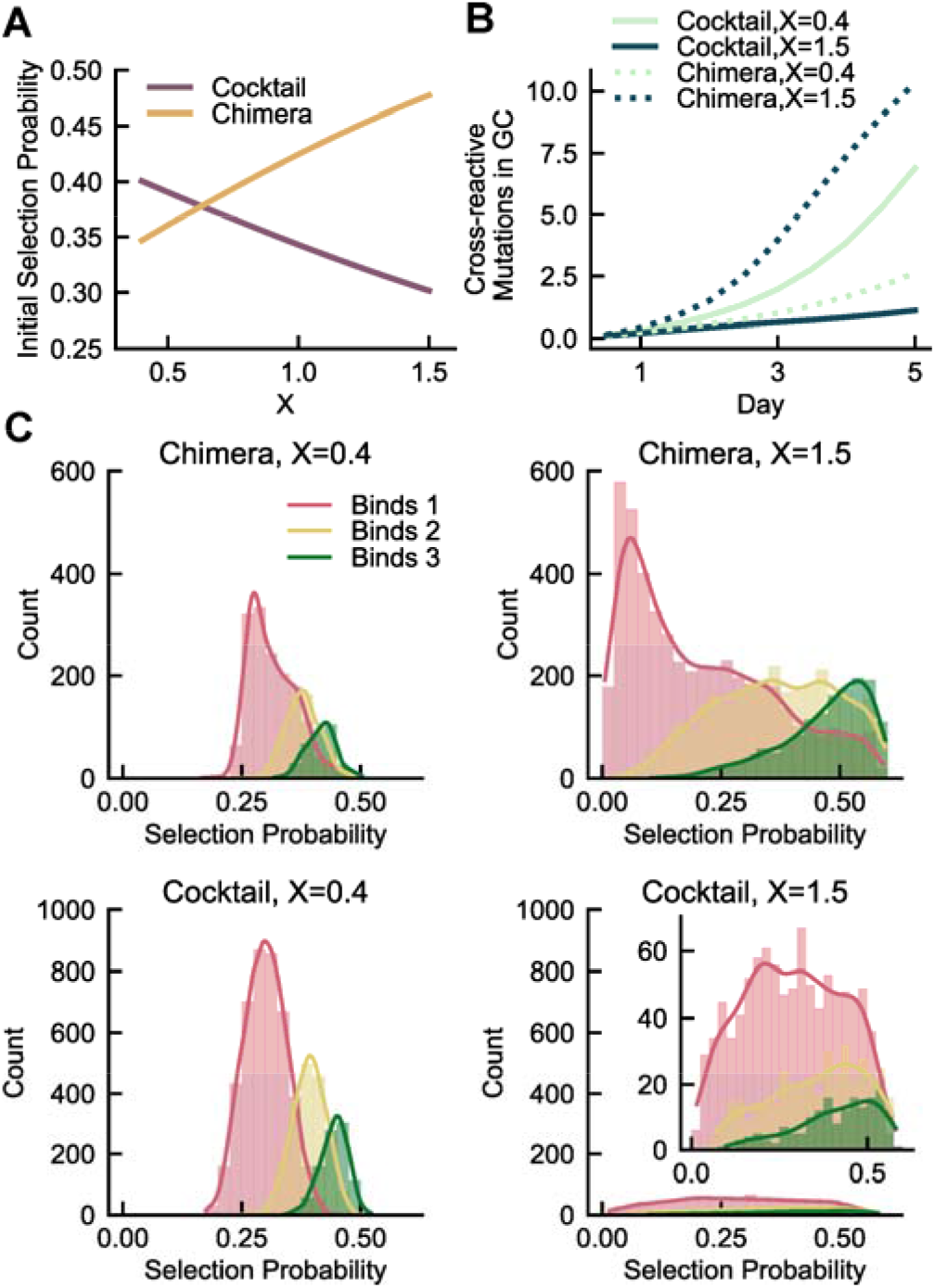
Mechanism of how T Cell selection stringency affects bnAb evolution. **(A)** Selection probability of the RBS-directed precursor at GC initiation as a function of T cell help stringency. **(B)** Average number of unique mutations found by RBS-directed B cells that increase affinity towards at least two variants in the first 5 days of affinity maturation. In panels A and B, the effects of T cell selection stringency are opposite for cocktail and chimeric immunogens. **(C)** Positive selection probabilities of unique RBS-directed B cell mutants in day 5 GCs, simulated under either stringent or permissive T cell selection conditions. The mutants are classified based on how many variants they can bind (from one to three). See also **Figure S3**.

For immunization with the cocktail immunogens, the difference in the amounts of antigen captured by cross-reactive and strain-specific B cells is small in the early GC when antigen is not limiting (**Fig. 1**). Therefore, increasing the stringency of how positive selection probability depends on the amount of antigen captured will not favor the bnAb precursors. The predominant difference between the cross-reactive and strain-specific B cells in the early GC is that only the former can capture diverse types of antigens so it can be positively selected by T cells with diverse epitope specificities, while the latter seek help from only a part of the repertoire of T helper cells. Cross-reactive B cells are promoted if this difference helps them during T cell selection. If selection stringency is permissive, each encounter with a cognate helper T cell will give similar chance of receiving positive selection signals. Cross-reactive B cells will encounter cognate T cells more frequently by capturing diverse epitopes, and despite the lower pMHC density of each epitope, the total probability of receiving help will be greater than strain-specific B cells that capture a similar total amount of antigen. Mathematical analysis of Eq. 9 (STAR Methods) shows that this is true when *x* < 1. Consistent with this analysis, our simulation results show that the selection probability of bnAb precursors in the early GC increases with decreasing *x* (**Fig. 4A**). The enhanced early selection probability allows bnAb precursors to more readily evolve future cross-reactive mutations (**Fig. 4B**). The cross-reactive mutants have distinct advantage over strain-specific mutants when selection is permissive and therefore selectively accumulate, but not when selection is stringent (**Fig. 4C**, bottom row). This is because, for less stringent selection, the ability of cross-reactive B cells to get positively selected by interacting with diverse T cells is amplified.

## DISCUSSION

Eliciting bnAbs is a necessary step towards a universal influenza vaccine that confers protection against seasonal variants and pandemic-causing novel strains. Most efforts to achieve this goal have focused on bnAbs that target a conserved region on the HA stem (Amitai et al., 2020; Impagliazzo et al., 2015; Lingwood et al., 2012; Sagawa et al., 1996). In this paper, we study the evolution of cross-reactive B cells that target the conserved HA RBS upon immunization with either a heterotrimeric RBS-enriched chimera or a homotrimeric cocktail of three antigenically distinct rsHAs (Caradonna et al., 2022).

Toward this end, we developed a computational model of affinity maturation upon vaccination with chimeric and cocktail immunogens. Our analyses of the pertinent processes and results (**Figs. 1-4**) provide new mechanistic insights into the factors that influence antibody repertoire development upon vaccination with different types of immunogens. We identify two important variables: the valency with which the antigen is bound to BCR, and the diversity of antigens captured by B cells. If bnAb precursors engage antigen multivalently, and strain-specific B cells cannot, as is true for the chimeric antigen (**Fig. 1**), then stringent selection of GC B cells by T helper cells promotes cross-reactive B cell evolution (**Figs. 3 and 4**). If the diversity of antigens captured is the principal difference between bnAb precursors and strain-specific B cells, as is true for the cocktail (**Fig. 1, 2**), then selection stringency must be permissive to promote bnAb evolution (**Figs. 3 and 4**). Because cross-reactive B cells are enriched in mice immunized with the chimeric immunogen we conclude that positive selection of B cells by T helper cells is a stringent constraint during GC reactions. Thus, our studies provide fundamental mechanistic insights on the role of T cell help during affinity maturation (Finney et al., 2018; Kuraoka et al., 2016; Mesin et al., 2016; Tas et al., 2016; Yeh et al., 2018), which will help improve vaccine design.

Our result suggests that one promising future direction would be to further optimize antigen valency using nanoparticles and epitope enrichment, to maximize the difference between the antigen capture capabilities of cross-reactive and strain-specific B cells. Alternatively, our model predicts that if T cell selection is permissive, a cocktail of antigens with distinct T cell epitopes can be highly effective at eliciting bnAbs. Is the stringency of T cell selection constant in the GC, or can it be dynamically regulated? There is evidence that some T follicular helper (Tfh) cells are of higher “quality” than others, that is, they can maintain a greater GC B cell/Tfh cell ratio (Havenar-daughton et al., 2017; Locci et al., 2013). Is it possible that the high-quality T cells are less discriminative? Moreover, we may also ask whether selection stringency can be controlled. Direct modulation of Tfh cell-B cell interaction by upregulating key surface molecules, such as SAP and SLAM has been suggested to make Tfh cells more potent helpers (Hu et al., 2013). Can such modifications and T cell depletion affect the T cell help stringency, and thus outcomes of vaccination using different antigens? Testing these hypotheses using our computational model and further experiments with designed immunogens will shed new light on basic questions in immunology and vaccine design.

## AUTHOR CONTRIBUTIONS

LY, TC, AKC and AGS designed research, LY carried out the calculations, AKC and LY analyzed the data, AKC, LY, TC, and AGS connected the experimental and computational results and wrote the paper.

## ACKNOWLEDGEMENTS

We thank Dr. Assaf Amitai for helpful discussions and critical insights. LY and AKC were supported by NIH grant U19AI057229 and funds from the Ragon Institute, and we acknowledge support from the NIH for R01AI146779 (A.G.S), P01AI89618-A1 (A.G.S) and T32 GM007753 (T.M.C.). LY also acknowledges partial funding from Kwanjeong Educational Foundation.

## STAR METHODS

### RESOURCE AVAILABILITY

○ Lead Contact
○ Materials availability
○ Data and code availability

### METHODS DETAILS

○ Affinity maturation simulation algorithm
○ Simulation of antigen capture
○ Selection by T cells
○ T cell epitope prediction and comparison

### RESOURCE AVAILABILITY

#### Lead contact

Further information and requests for resources and reagents should be directed to and will be fulfilled by the lead contact, Arup Chakraborty.

#### Materials availability

This study did not generate new unique reagents.

#### Data and code availability

- Simulation data have been deposited on Mendeley at doi: 10.17632/2kt95vthcs.1 and are publicly available.
- All original code has been deposited on Mendeley at doi: 10.17632/2kt95vthcs.1 and are publicly available.
- Any additional information required to reanalyze the data reported in this paper is available from the lead contact upon request.

## METHOD DETAILS

### Affinity maturation simulation algorithm

As described in the main text, we simulate in-silico germinal centers in which B cells capture antigen and then compete for help by T cells in each cycle, for 28 cycles. The stochastic GC simulation is repeated 1,000 times. We keep track of the following quantities: the number of GC B cells that target each epitope (rsH3, rsH4, or rsH14 off-target B cells or RBS-directed B cells), the binding affinities of the GC B cells, the mutations that are carried by the RBS-directed B cells, and the probabilities of positive selection of RBS-directed B cells at each round. For reporting the RBS-directed B cell fractions (see Figure 3), all B cells from the 1,000 GCs are first pooled together, and then the fraction is calculated.

The amounts and types of antigens captured by the B cells are determined by simulating the immunological synapse between the B cell and the FDC. BCRs first cluster with antigens, then internalize them by applying force (see sections **Model Development** and **Antigen capture depends on antigen design and cross-reactivity of B cells** in the main text). Then, the probability of positive selection by T cells is determined based on the amount and diversity of the antigens captured by the B cell, relative to other competing B cells (see Eq. 9 and associated description in the main text). We provide further detail and analyses of these steps below.

### Simulation of antigen capture

The immunological synapse is modeled as a circle of radius 0.5 µm divided up into lattice points with an interval of 10 nm that can be occupied by the antigens and BCRs. No two homotypic molecules are allowed on the same lattice site, but a BCR and an Ag molecule can occupy the same site. To begin the simulation, 120 BCRs and 120 Ag molecules are randomly distributed on the lattice sites. During the clustering phase, BCR and Ag molecules diffuse freely. In each time step, each molecule randomly chooses one of the four neighboring sites, then move to it with the probability of,

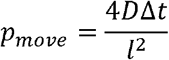

where *D* =5 ×10^4^ *nm*^2^s^-1^ is the diffusion constant for both Ag and BCR (Fleire et al., 2006b), and l= 10 nm is the lattice size. For clusters of BCRs and Ags, only those containing up to 3 molecules are allowed to diffuse and the diffusion coefficient is reduced to *D/M* where *M* is the number of molecules in the cluster (Meakin, 1984). The move is completed if the new sites are not blocked for the diffusing molecules. If any of the new sites are already occupied or are outside the boundary of the immunological synapse, the move is not accepted and the simulation continues to the next step.

When the distance between a BCR and an Ag molecule is either 0 or 1 lattice site, they can bind, as described in the main text. When several free epitopes on the Ag molecules are cognate for the BCR, one is randomly chosen upon binding. The sizes of clusters stabilize within a few seconds (data not shown), so we simulate the clustering phase for 10 seconds.

When the extraction phase begins, BCRs and any Ag molecules bound to them stop diffusing, but free Ag molecules still diffuse. A pulling force is applied to each BCR, which affects the antigen extraction as described in the main text (see Equations 6-8). The simulation terminates once all BCRs are internalized, and the number and types of internalized Ag molecules are calculated.

The simulation of antigen capture is computationally intensive, so repeating it for thousands of B cells for each round of affinity maturation is impractical. Therefore, we first run the antigen capture simulations to determine the mapping between the binding affinities of a B cell and the amount of antigen it captures, then use this mapping to quickly determine how much antigen each B cell captures during the affinity maturation simulations. To obtain the mapping for a strain-specific B cell, we run 30 independent simulations of antigen capture for each value of binding affinity between −13.8 and −20.8 *k*_*B*_*T* with an maturation simulations. To obtain the mapping for a strain-specific B cell, we run 30 independent interval of 0.5 *k*_*B*_*T*. The mean amount of antigen captured is determined at each point. This affinity range covers the limits of B cell affinities relevant in our affinity maturation simulation. The amount of antigen captured by a B cell is determined from standard linear interpolation using the two nearest points to its binding affinity. For the RBS-directed B cells, we run the antigen capture simulations for a set of grid points on a three-dimensional grid, where each axis corresponds to the binding affinity towards one variant, ranging between −13.8 and −20.8 *k*_*B*_*T* with an interval of 0.5 *k*_*B*_*T*. The amount of antigen points on a three-dimensional grid, where each axis corresponds to the binding affinity towards one captured by a given B cell is obtained from a standard trilinear interpolation using the eight nearest points.

### Selection by T cell

The main text describes how the probability of positive selection by T cells depends on both the amount and the diversity of the captured antigens (see Equation 9). Here, we provide a mathematical analysis of why immunization with the cocktail antigen favors cross-reactive B cells in the early GC when T cell help is permissive, but not when it is stringent (see Figure 4).

The bnAb precursors and off-target B cells with low affinities capture similar amounts of total antigen.

For simplicity, let us assume that the amounts of antigen captured are equal. That is, *A*_1_+ *A*_2_+ *A*_3_+ = *A*_1_+_*off*_ where *A*_1_+ *A*_2_+ *A*_3_+ >0 are the amounts of the three variants captured by an RBS-directed B cell, and *A*_1_+_*off*_ is the amount captured by an off-target B cell that, without loss of generality, is assumed to target only the first variant.

Eq. 9 is a monotonically increasing function of the numerator 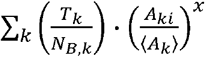. Therefore, tounderstand how the positive selection probability of the RBS-directed B cell compares with that of the off-target B cell, we will compare q_RBs_, defined as 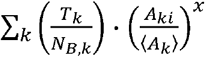 and q_*off*_, defined as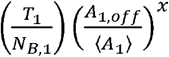

In our model, we assume that equal numbers of T cells target the epitopes from each of the three rsHA variants. That is, *T*_1_ =*T*_2_ =*T*_3_Also, each off-target B cell is randomly assigned the target variant with equal probability. Since there is a relatively large number of founder B cells (99 off-target B cells), we can approximate that the number of B cells that capture each variant are equal, i.e. *N* _*B*,1_ = *N* _*B*,2_ = *N* _*B*,3_=

Also, the mean amount of antigen internalized by B cells that recognize antigen, *k*, ⟨*A*_*k*_⟩is equal for all at the beginning of the GC because all B cells have the same affinity. Since GCs contain thousands of B cells, this equality also holds well due to symmetry even when B cells begin to mutate, at least in early GCs. Taken together, the following equality holds.

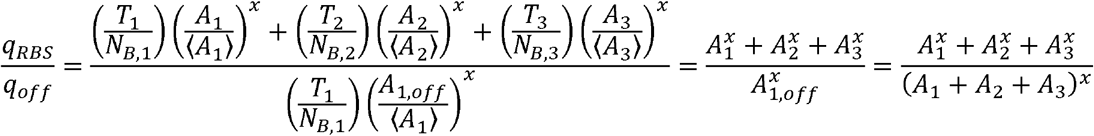

To show that the RBS-directed B cells are favored for positive selection when T cell help is permissive, we will prove the following inequality:

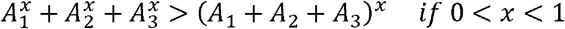

Consider the function 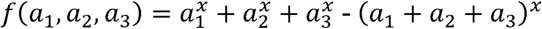defined for positive real numbers *a*_1_, *a*_2_, *a*_3_. The partial derivatives are always positive if 0 < *x* <1:

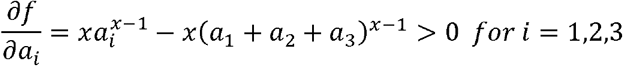

because and *a*_*i*_ < *a*_1_ + *a*_2_ + *a*_3_ *x* − 1 < 0.

Assume that there exists *A*_1_, *A*_2_, *A*_3_ >0 such that *f* (*A*_1_, *A*_2_, *A*_3_)=*S* ≤0. Then, for any *a*_1,_ *a*_2_, *a*_3_, such that *a*_1_ ∈ (0,*A*_1_), *a*_2_ ∈ (0,*A*_2_), *a*_3_ ∈ (0, *A*_3_) the following inequality must be true.

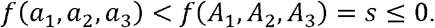

However, *f* is a continuous function and lim_al → 0_,a2_→_,a3_→_ *f* (*a*_1_, *a*_2_, *a*_3_) =0. Therefore, there must exist δ > 0 such that

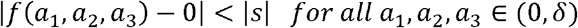

which is contradictory.

Therefore, *q*_*RBS*_ > *q*_*Off*_ when *x* < 1, and by simple extension, 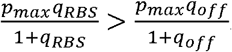. That is, despitecapturing the same amount of antigen, the RBS-directed B cell has a higher probability of positive selection because of capturing diverse T cell epitopes. By similar analysis, it can be shown that when *x* > 1 the opposite is true, and the RBS-directed B cells have a lower probability of positive selection.

### T cell epitope prediction and comparison

T cell epitopes in the rsH3, rsH4, and rsH14 antigens (Figure 2) as well as in the rabbit serum albumin and human serum albumin (Fig. S2) are predicted with IEDB MHCII binding prediction tool. The following settings are used: Prediction Method – IEDB recommended 2.22; Select species/locus – mouse, H-2-I; Select MHC allele – H2-I-A^b^; Select length – 15. The predicted peptides are sorted by the percentile rank given by the IEDB tool, and the peptides in the top 20 percentile are chosen for the pairwise comparisons of the epitopes in different variants. This value corresponds to roughly 3000 nM predicted half-maximal inhibitory concentration (IC50) value. We choose this cutoff to be comprehensive because most immunogenic MHC II T cell epitopes have an IC50 value under 1,000 nM (Southwood et al., 1998). The 9-mer cores associated with the peptides, predicted by the smm_align method, are then used for the pairwise comparisons.

