## Supplemental Figures and Table for "Mechanisms that promote the evolution of cross-reactive antibodies upon vaccination with designed influenza immunogens"

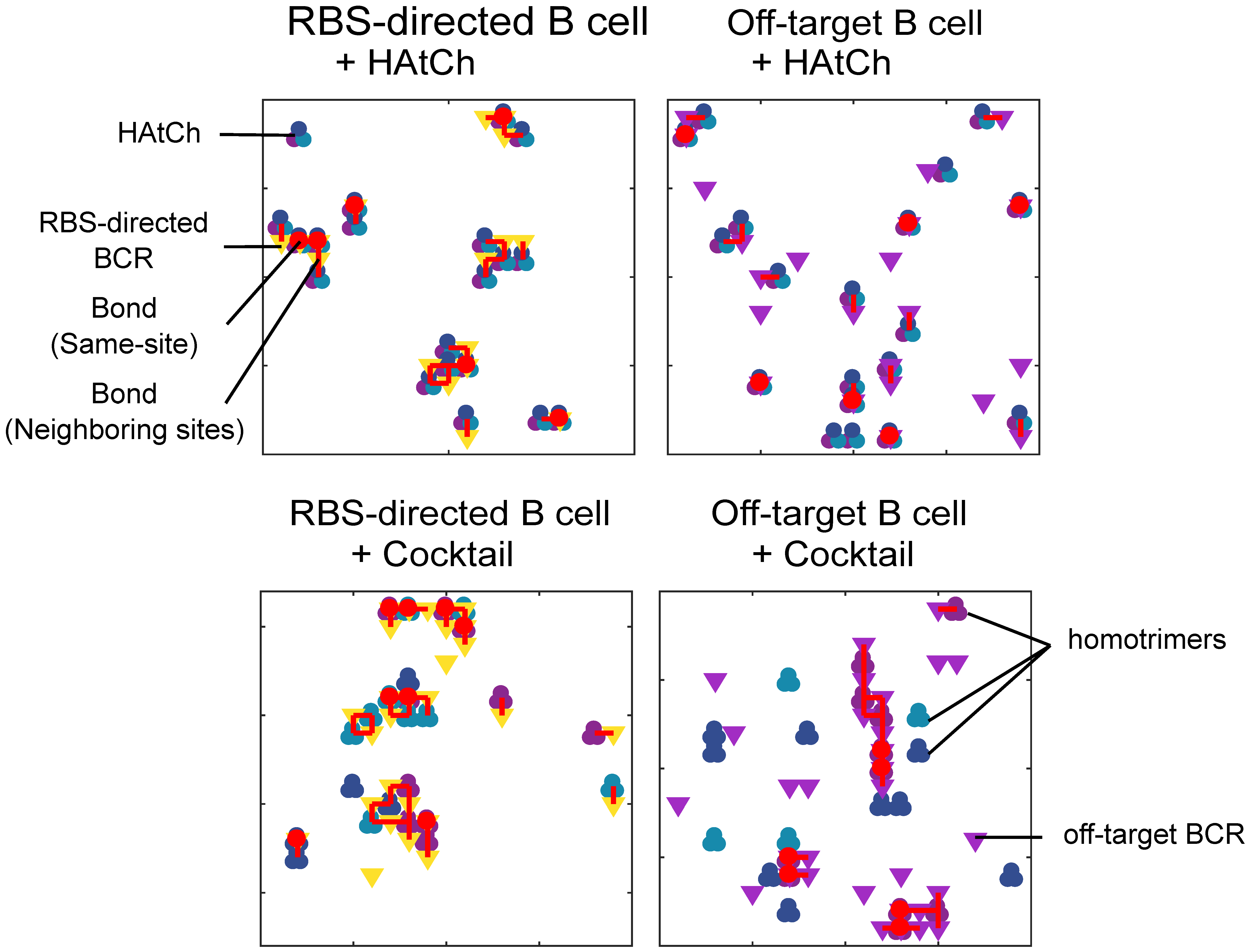

**Figure S1. Related to Figure 1. Clustering of antigens and BCRs in simulations of antigen capture by B cells**. Examples of the immunological synapses from the simulations are shown. The simulation recapitulates how antigen design and B cell cross-reactivity affect the clustering of BCRs and antigens, as described in Figure 1A. A single HAtCh molecule binds to several BCRs of an RBS-directed B cell, but only one BCR of an off-target B cell at most. All three types of homotrimers in the cocktail cluster with the BCRs of an RBS-directed B cell, but only the target variant cluster with the BCRs of an off-target B cell.

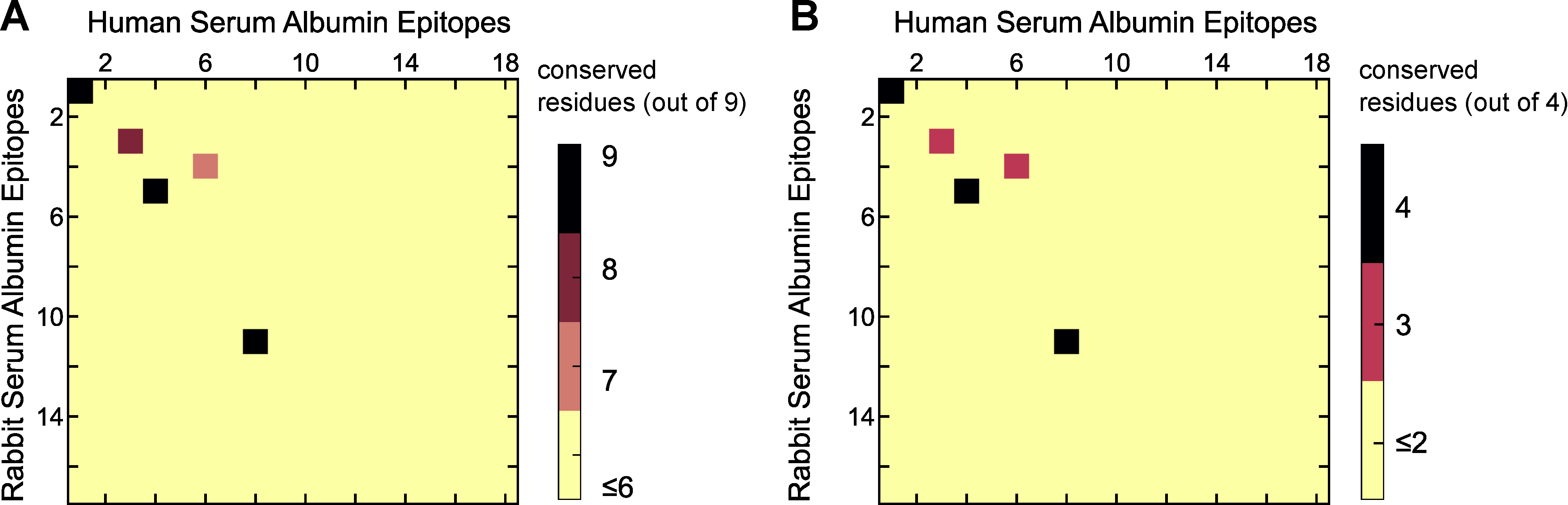

**Figure S2. Related to Figure 2. T Cell Epitope Analysis of Rabbit and Human Serum Albumin. (A)** Number of conserved residues in pairwise comparison of the 9-mer cores of the top 20 percentile predicted 15-mer T cell epitopes in rabbit and human serum albumin. **(B)** Number of conserved residues in pairwise comparisons focusing on the P2, P5, P7, P8 residues of the 9-mer cores. Each axis corresponds to the ranks of the top 20 percentile predicted 15-mer T cell epitopes

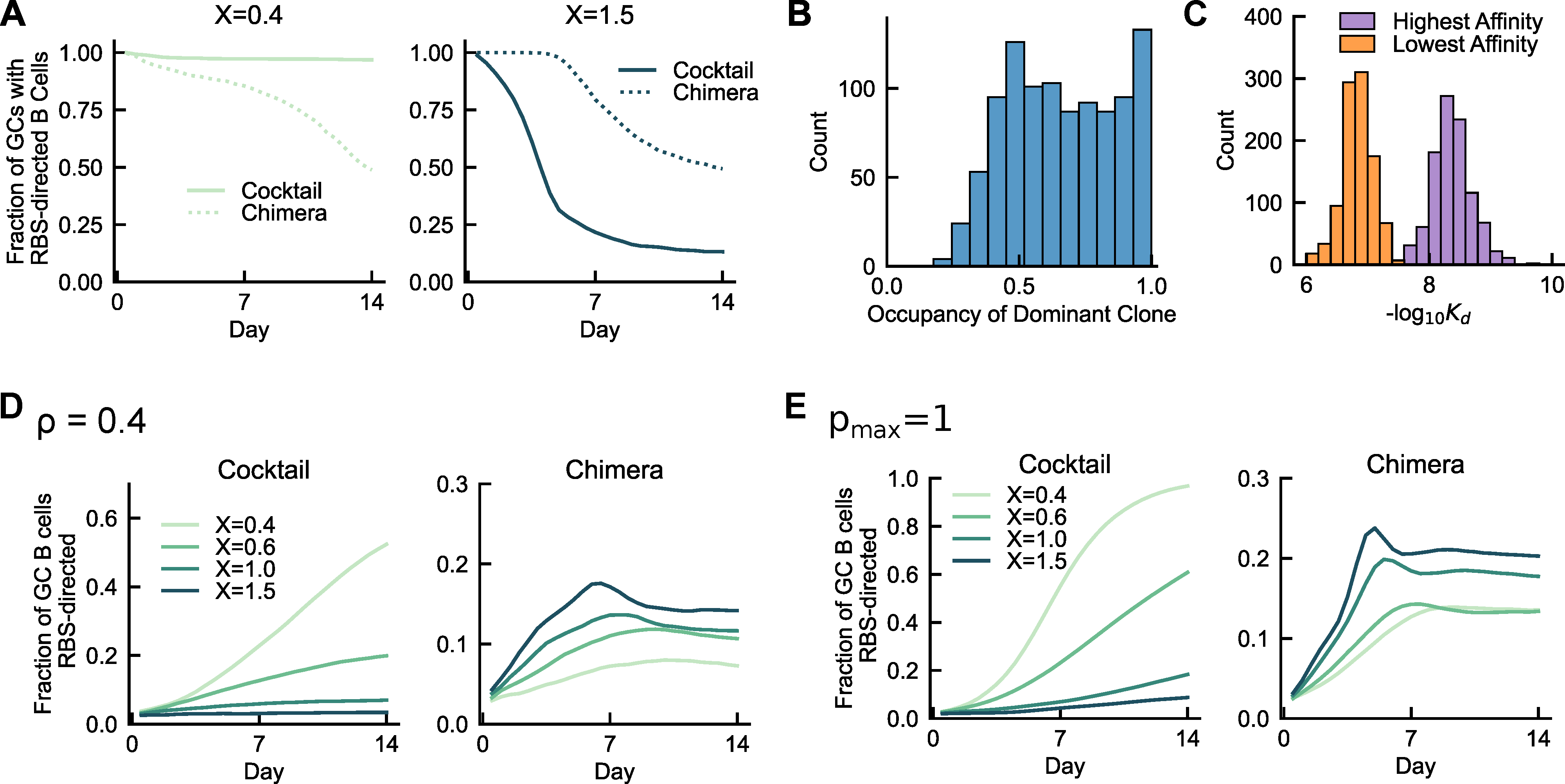

**Figure S3. Related to Figure 3. Evolution of RBS-directed B cells in Germinal Centers. (A)** The fraction of GCs that contain at least one RBS-directed B cell. When selection by T cells is permissive, RBS-directed B cells survive in most GCs after immunization with the cocktail, while they slowly become extinct in an increasing number of GCs after immunization with the chimeric antigen. When selection is stringent, RBS-directed B cells rapidly become extinct in many GCs after immunization with the cocktail. They survive in most GCs after immunization with the chimeric antigen for the first several days, then, since affinity increases quickly and the advantage of cross-reactivity decreases in late GC, some GCs lose the RBS-directed B cells. **(B-C)** Heterogeneities among 1000 GC simulations on day 9, when T cell selection is stringent (*x* = 1.5), resulting from stochasticity in evolution. (B) Fraction of GC B cells that belong to the most dominant clonal lineage. The GCs show varying degrees of homogenization. (C) The highest and lowest affinity of positively selected B cells on day 9 from each GC. There is a ~10 fold affinity difference among positively selected B cells within the same GCs, which is highlighted by the difference between the two peaks. Across all the GCs, B cells with an even wider range of affinities coexist. **(D-E)** The fraction of GC B cells that are RBS-directed when parameters *ρ* and *p_max_* are changed, showing the same qualitative trend as the main result in Figure 3. (D) *ρ* is 0.4, instead of 0.7. (E) *p_max_* is 1, instead of 0.6.

**Table S1. Related to Figure 2. MHC II Binding T Cell Epitopes Predicted by Computational Tools.** The 15-mer epitopes are grouped based on the 9-mer cores. Peptides in top 20 percentile rank are shown, corresponding to roughly 3000 nM predicted half maximal inhibitory concentration (IC50) value. The prediction was made by MHCII binding prediction tool using IEDB recommended 2.22 method, for the mouse MHC H2-IAb allele. See STAR METHODS for details.

| **rsH3** | | | **rsH4** | | | **rsH14** | | |
| --- | --- | --- | --- | --- | --- | --- | --- | --- |
| **Core** | **Peptide** | **Rank** | **Core** | **Peptide** | **Rank** | **Core** | **Peptide** | **Rank** |
| FNWTGVTQN | NNESFNWTGVTQNGV | 1.6 | FQWSTVTQN | IAEEFQWSTVTQNGV | 2.8 | FTWNGVTVD | IAEQFTWNGVTVDGV | 5.85 |
|  | NESFNWTGVTQNGVS | 1.65 |  | FIAEEFQWSTVTQNG | 2.9 |  | AEQFTWNGVTVDGVS | 6.15 |
|  | ESFNWTGVTQNGVSA | 1.7 |  | AEEFQWSTVTQNGVS | 3.1 |  | EQFTWNGVTVDGVSA | 6.4 |
|  | FNNESFNWTGVTQNG | 1.7 |  | EEFQWSTVTQNGVSA | 3.2 |  | FIAEQFTWNGVTVDG | 7.1 |
|  | SFNWTGVTQNGVSAS | 3.4 |  | EFQWSTVTQNGVSAS | 4.25 |  | QFTWNGVTVDGVSAS | 9.9 |
|  | FNWTGVTQNGVSASC | 4.9 | EEFQWSTVT | EFIAEEFQWSTVTQN | 3 | EQFTWNGVT | EFIAEQFTWNGVTVD | 7.05 |
| ESFNWTGVT | EFNNESFNWTGVTQN | 1.8 | VSASCSRAN | TQNGVSASCSRANVS | 3.35 | YVKQGSLML | CPKYVKQGSLMLATG | 9.75 |
| ITPNGSIPN | KSECITPNGSIPNDK | 6.75 |  | QNGVSASCSRANVSD | 3.75 |  | NCPKYVKQGSLMLAT | 10.4 |
|  | CKSECITPNGSIPND | 8.2 |  | VTQNGVSASCSRANV | 4.15 |  | PKYVKQGSLMLATGG | 10.9 |
|  | SECITPNGSIPNDKP | 8.25 |  | NGVSASCSRANVSDF | 4.5 |  | KYVKQGSLMLATGGA | 15.5 |
|  | ECITPNGSIPNDKPF | 9.25 |  | GVSASCSRANVSDFF | 9.05 |  | GNCPKYVKQGSLMLA | 15.8 |
|  | CITPNGSIPNDKPFQ | 13.7 |  | VSASCSRANVSDFFN | 13 |  | YVKQGSLMLATGGAG | 19.5 |
|  | ITPNGSIPNDKPFQN | 16 | VTQNGVSAS | TVTQNGVSASCSRAN | 4.05 | KNGLYGPIN | TGKNGLYGPINVTKE | 10.2 |
| YASLRSLVA | DVPDYASLRSLVASS | 6.8 |  | FQWSTVTQNGVSASC | 5.45 |  | LTGKNGLYGPINVTK | 14 |
|  | VPDYASLRSLVASSG | 6.8 |  | STVTQNGVSASCSRA | 7.3 |  | WLTGKNGLYGPINVT | 16.5 |
|  | YDVPDYASLRSLVAS | 7.2 |  | WSTVTQNGVSASCSR | 8.55 | YLWGVHHPS | VRLYLWGVHHPSNIG | 10.5 |
|  | PDYASLRSLVASSGT | 7.3 |  | QWSTVTQNGVSASCS | 9.85 |  | RLYLWGVHHPSNIGD | 11.1 |
|  | YASLRSLVASSGTLE | 9.3 | LNTAVPIGS | STILNTAVPIGSCVS | 8.7 |  | YVRLYLWGVHHPSNI | 11.4 |
|  | DYASLRSLVASSGTL | 9.4 | AVPIGSCVS | TILNTAVPIGSCVSK | 9.25 |  | LYLWGVHHPSNIGDQ | 14 |
| VTQNGVSAS | WTGVTQNGVSASCSR | 8.05 |  | ILNTAVPIGSCVSKC | 11.5 |  | YLWGVHHPSNIGDQR | 16.3 |
|  | TGVTQNGVSASCSRR | 8.1 |  | LNTAVPIGSCVSKCH | 13.5 |  | SYVRLYLWGVHHPSN | 17.5 |
|  | NWTGVTQNGVSASCS | 9.25 | WGVHHPPNI | ARLYIWGVHHPPNIG | 9.3 | YGPINVTKE | GKNGLYGPINVTKEN | 11.9 |
|  | GVTQNGVSASCSRRS | 12.6 |  | RLYIWGVHHPPNIGD | 10.4 |  | KNGLYGPINVTKENT | 12.7 |
|  | VTQNGVSASCSRRSS | 14.5 |  | LYIWGVHHPPNIGDQ | 11.6 |  | NGLYGPINVTKENTG | 19.5 |
| SECITPNGS | KCKSECITPNGSIPN | 8.15 |  | YIWGVHHPPNIGDQR | 13.4 | VTVDGVSAS | FTWNGVTVDGVSASC | 14 |
| PDYASLRSL | PYDVPDYASLRSLVA | 8.55 | TILNTAVPI | QKKSTILNTAVPIGS | 11 | FNSIGNLIA | IIFNSIGNLIAPRGH | 15.5 |
| KNGLYPALN | TGKNGLYPALNVTMP | 8.85 |  | KSTILNTAVPIGSCV | 11.7 |  | SIIFNSIGNLIAPRG | 15.5 |
|  | LTGKNGLYPALNVTM | 9.8 |  | KKSTILNTAVPIGSC | 11.9 |  | DSIIFNSIGNLIAPR | 16 |
|  | WLTGKNGLYPALNVT | 11 |  | SQKKSTILNTAVPIG | 15 |  | GDSIIFNSIGNLIAP | 16.5 |
| GLYPALNVT | GKNGLYPALNVTMPN | 10.6 | YIWGVHHPP | YARLYIWGVHHPPNI | 13.5 | SIIFNSIGN | PGDSIIFNSIGNLIA | 16.5 |
|  | KNGLYPALNVTMPNK | 10.8 | IFIERPTAV | EWDIFIERPTAVDTC | 15.5 | VFIERPTAM | TWDVFIERPTAMDTC | 16.5 |
| YPALNVTMP | NGLYPALNVTMPNKE | 11 |  | WDIFIERPTAVDTCY | 16 |  | TTWDVFIERPTAMDT | 17 |
| LNVTMPNKE | GLYPALNVTMPNKEQ | 12.5 |  | AEWDIFIERPTAVDT | 16.5 |  | WDVFIERPTAMDTCY | 17.5 |
|  | LYPALNVTMPNKEQF | 18 |  | GAEWDIFIERPTAVD | 17.5 |  | DTTWDVFIERPTAMD | 18.5 |
| SLVASSGTL | ASLRSLVASSGTLEF | 15.5 | KKSTILNTA | NSQKKSTILNTAVPI | 17 | YQSLRSILA | PDYQSLRSILASSGS | 17 |
|  | SLRSLVASSGTLEFN | 17.5 | YQSLRSILA | DVPDYQSLRSILANN | 17.5 |  | DVPDYQSLRSILASS | 17.5 |
|  | LRSLVASSGTLEFNN | 19 |  | FDVPDYQSLRSILAN | 17.5 |  | VPDYQSLRSILASSG | 17.5 |
| FKIRSGKSS | RGYFKIRSGKSSIMR | 17.5 |  | VPDYQSLRSILANNG | 17.5 |  | FDVPDYQSLRSILAS | 18 |
|  | PRGYFKIRSGKSSIM | 18 |  | PDYQSLRSILANNGK | 19 |  | DYQSLRSILASSGSL | 18.5 |
|  | GYFKIRSGKSSIMRS | 19.5 |  |  |  | IYWTLVNPG | ISIYWTLVNPGDSII | 17.5 |
|  |  |  |  |  |  |  | SIYWTLVNPGDSIIF | 19.5 |
|  |  |  |  |  |  | RLYLWGVHH | GSYVRLYLWGVHHPS | 18.6 |
|  |  |  |  |  |  | SILASSGSL | QSLRSILASSGSLEF | 19 |
|  |  |  |  |  |  |  | YQSLRSILASSGSLE | 19 |
|  |  |  |  |  |  | KYVKQGSLM | IGNCPKYVKQGSLML | 19.4 |
